# Single-Cell Signature Explorer for comprehensive visualization of single cell signatures across scRNA-seq data sets

**DOI:** 10.1101/621805

**Authors:** Frédéric Pont, Marie Tosolini, Jean Jacques Fournié

## Abstract

The momentum of scRNA-seq data sets prompts for simple and powerful tools exploring their meaningful signatures. Here we present Single-Cell_Signature_Explorer (https://sites.google.com/site/fredsoftwares/products/single-cell-signature-explorer), the first method for high throughput scoring at single cell level of any gene set-based signature and visualization across t-SNE. By scanning data sets for single or combined signatures, it quantitatively and qualitatively maps any multi-gene feature, exemplified here with signatures of cell lineages, biological hallmarks and metabolic pathways in large scRNAseq datasets of human PBMC, lung cancer and adult testis.

## INTRODUCTION

The development of single cell RNA sequencing (scRNA-seq) techniques yields an increasing number of data sets available to the scientific community. These comprise reference samples of human tissues and organs from healthy individuals as in the Human Cell Atlas repository (1), or published studies of samples from dysfunctional or pathological tissues. Therefore, it becomes important to assess in each single cell from such large data sets any kind of hallmark possibly defined by a gene set, such as cell lineage, maturation, biological and metabolic signatures, and visualize this feature across the t-distributed stochastic neighborhood embedding (t-SNE) plots (2) in a global and comprehensive viewpoint. This is conceptually similar to gene set enrichment analysis (GSEA) (3), which finds functional significance of genes differentially expressed between bulk transcriptomes of two series of samples. Currently however, there is no method for performing this at a single cell level and comparatively to all the data set’s cells, in a reasonable time. Here we report Single Cell Signature Explorer, an algorithm which scores rapidly many gene set-defined signatures at the single cell level and allows their straighforward and interactive visualization across t-SNE plots of entire scRNA-seq data sets, together with highly informative examples of its application.

## MATERIALS AND METHODS

### Single Cell experiments

Data for 4k and 8k PBMC obtained by 10x Genomics 3’ chemistry V2 were downloaded from 10x Genomics website (https://support.10xgenomics.com/single-cell-gene-expression/datasets/2.1.0/pbmc8k, https://support.10xgenomics.com/single-cell-gene-expression/datasets/2.1.0/pbmc4k).

Data for 19k of lung tumor and normal adjacent cells (4) obtained by 10x Genomics 3’ chemistry V2 were downloaded from Array Express E-MAAB-6149 and E-MTAB-6653.

The human spermatogenesis data (5) obtained by 10x Genomics 3’ chemistry V2 were downloaded from GEO data set GSE120508. The 60k melanoma data set obtained by MARS platform (6) were downloaded from GEO data set GSE12313.

### t-SNE maps computation

The t-SNE map were produced by Seurat (7) as follows: Row data were computed with CellRanger 2.2.0 and then loaded in a R session with the Seurat package. Samples were individually filtered using UMI and percentage of mitochondrial genes criteria. Samples were then merged using correction to align data sets as described in (7). t-SNE coordinates were then calculated using the 11 first PCA and exported in a table.

### Noise reduction of data sets

Single cell transcriptomic technology is powerful to explore gene expression at the single cell level, but may yield scRNAseq datasets in which technical noise can hinder biological variability. This technical noise may arise from both gene sampling fluctuations and cell-to-cell variations in sequencing efficiency (8). To reduce such technical noise from raw UMI data sets, we applied scRNAseq DataSet Noise Reductor (https://sites.google.com/site/fredsoftwares/products/single-cell-signature-explorer). Briefly, this algorithm successively applies median normalisation and Freeman-Tuckey transform to the raw UMI count of each gene and single cell from any data set such as to stabilize its technical noise at a comparable level between genes and across cells (9). A detailed description of scRNAseq DataSet Noise Reductor results is provided in the supplemental methods. This section shows that Single-Cell Signature Explorer indifferently processes scRNAseq datasets of either raw UMI data, noise-reduced UMI data, or UMI data normalized by Seurat. Noise reduction is optional, under the responsibility of users and in any case a preliminary to Single-Cell Signature Explorer. Whatever the signature, Single-Cell Signature Explorer produces similar t-SNE images from these various types of UMI data, but requires considerably longer computing time for the noise-reduced data.

### Single-Cell Signature Explorer

Single-Cell Signature Explorer is a package of four successive tools dedicated to high throughput signature exploration in single-cell analysis:

1. Single-Cell Signature Scorer computes for each cell a signature score.
2. Single-Cell Signature Merger collates the signature scores table with t-SNE coordinates.
3. Single-Cell Signature Viewer displays signatures scores on a t-SNE map.
4. Single-Cell Signature Combiner displays arithmetic combinations of two signatures scores on a t-SNE map.

These four softwares have been developed with usability, low memory usage and performances in mind. They require no complex command lines or computing skills, they can be used on a laptop but also scale up very well on powerful workstations for fast computations: results are obtained within seconds in function of the gene set size.

#### Single-Cell Signature Scorer

This algorithm computes in each single cell transcriptome a score for any each gene set from a database. More than 17,000 curated gene sets from MSigDB(3, 10) and Reactome (11), as well as additional user-defined gene sets can be computed by the software. Each gene set is a list of HUGO gene symbols in a text file format labeled by the name of the gene set. The database can also be implemented by additional custom pathways composed of text files for user-defined list of HUGO gene symbols. Scores were calculated as follows: The score of the gene set *GS_x_* in cell *C_j_* was computed as the sum of all UMI for all the *GS_x_* genes expressed in *C_j_*, divided by the sum of all UMI expressed by *C_j_*:

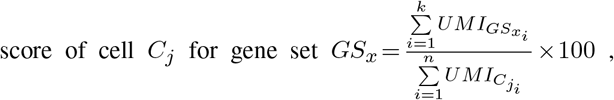

where *UMI_GS_x_i___* are the UMI of the *k* genes from *GS_x_* found in the cell *C_j_*. We also implemented the possibility to subtract the contribution of a gene when it is preceded by a “−” sign in the gene list. Single-Cell Signature Scorer was developed in Go language v1.11.1 (https://golang.org/). The software is multi-threaded to take full advantage of multicore processors. Single-Cell Signature Scorer can compute 13 million scores (1000 gene sets for 13300) cells in less than 4’30” on a bi-Xeon E5-2687w-v3 workstation. Single-Cell Signature Scorer was compiled for GNU Linux and Microsoft© Windows™64 bits, and can be compiled for any platform using cross-compilation feature of Go.

#### Single-Cell Signature Merger

This very fast tool, written in Go (https://golang.org/), yields a table merging for each cell the signature score and the t-SNE coordinates. This software takes as input the score tables created by Single-Cell Signature Scorer and t-SNE coordinates produced by Seurat (7) to produce tables compatibles with the Single-Cell Signature Viewer (see below). This software is multi-threaded and can merge in parallel a set of score tables with a t-SNE coordinates table.

#### Single-Cell Signature Viewer

This tool was developed to display the gene set scores computed by Single-Cell Signature Scorer on a t-SNE map. It takes as input a table score merged with t-SNE coordinates. Single-Cell Signature Viewer was written in R (12) with the package Shiny (13) to draw in real time a colored score scale of signatures selected from a drop-down list on t-SNE maps. Since this scaling is sensitive to outliers, the viewer draws a density distribution of scores and provides a color scale cursor allowing users to prune such potential outliers.

#### Single-Cell Signature Combiner

This tool was developed to display the combination of two gene sets scores previously computed by Single-Cell Signature Scorer on a t-SNE map. User must select two signatures from two drop-down lists. Before combining the two corresponding score sets, they are normalized between [0-1] to be comparable. The operators available to compute the combination are [− +], thus user selects to add or subtract two signatures prior to visualize the resulting score on one t-SNE map. Single-Cell Signature Combiner takes as input a table score merged with t-SNE coordinates. Single-Cell Signature Combiner was written in R (12) with the package Shiny (13) to draw in real time on t-SNE maps a colored score scale of two signatures selected from the drop-down lists. As above, since this scaling is sensitive to potential outliers, the viewer draws a density distribution of scores and provides a color scale cursor allowing the user to prune such outliers.

#### Code availability

**Single-Cell Signature Explorer** was developed using the high performance cross-platform Go language v1.11.1 (https://golang.org/) and the t-SNE viewers were developed using the cross-platform R (12) language with the package Shiny (13). Files can be accessed at (https://sites.google.com/site/fredsoftwares/products/single-cell-signature-exp)

### Software Benchmarks

We compared Single-Cell Signature Explorer with AUCell (14), Seurat CellCycleScore (7), and GSVA/ssGSEA (15, 16). Their respective computation efficiency and display of the results on the t-SNE map were evaluated.

#### Signature Scores computation speed

Benchmarks were performed on a Linux Xubuntu 18.10 workstation with two processors Xeon E5-2687w-v3 and 128Go RAM. KEGG database was generated from MSigDB v6.2 (http://www.broad.mit.edu/gsea/msigdb/index.jsp). For Single-Cell Signature Scorer, KEGG database was downloaded from the software web site where additional, ready-to-use databases of 17,000 gene sets for Human, Mouse, Rat, Zebra Fish, and Macaccus, obtained from MSigDB v6.2 are also available (https://sites.google.com/site/fredsoftwares/products/databases). For the ssGSEA speed test, both software issues and the too long computation time (>12 hrs) for the entire KEGG database did not permit this evaluation. Hence this speed test was only performed on two gene sets and its result was extrapolated. Since Seurat function CellCycleScore can only compute gene sets with anti-correlated expression levels, Seurat computation time for all gene sets of KEGG database could not be performed.

## RESULTS AND DISCUSSION

Although GSEA measures the relative enrichment of functionally defined sets of genes, it cannot characterize the gene signature from an isolated transcriptome, while the related ssGSEA (15), GSVA (16) and AutoCompare_SES (17) achieve this by scoring the intrinsic enrichment of pre-defined gene sets in any single sample transcriptome. Nevertheless, both algorithms rely on statistics inappropriate for single cell transcriptomes mostly composed of zero values, and where the missing genes of each cell vary extensively across the entire scRNA-seq data set. To address these issues with minimal computing time, here we introduce Single-Cell Signature Explorer, a package of four softwares dedicated to high throughput signature exploration in single-cell transcriptome analysis. The score of the gene set GS_*i*_ in cell C_*j*_ is computed as the sum of all UMI for all the GS_*i*_ genes expressed in C_*j*_, divided by the sum of all UMI expressed by C_*j*_. Using this, the scores of each single cell for thousands gene sets (*e.g*. 17,000 gene sets from the Molecular Signatures Database (MSigDB) (http://www.broad.mit.edu/gsea/msigdb/index.jsp) are computed within few minutes, and these scores are then visualized across entire scRNA-seq data sets.

As a first example, the scRNA-seq datasets of 4k and 8k PBMC from one healthy donor were downloaded (https://support.10xgenomics.com/single-cell-gene-expression/datasets), integrated, processed and the resulting t-SNE was plotted. In parallel, each of these 12k single cells was individually scored for each of the 17,000 MSIgDB gene sets. As negative controls, these cells were also scored likewise for 1000 random gene sets, each composed of *n* randomly sampled genes without replacement (RGS1-RGS1000). The PBMC data set comprises a cluster of (*n*=1731) B lymphocytes which can be defined either as cells expressing the B cell-identifying *CD19* gene alone (*n*= 883 cells), or as (*n*= 1826) cells scoring for a ‘B cell’ signature composed of the four genes *CD79A, CD79B, CD19, MS4A11*. Both cell counts and ROC curves indicated that the B cell signature outperformed the single gene in spotting all B lymphocytes from the data set, including the few (*n*= 109) plasmocytes which mapped outside of the B cell cluster (**Figure 1**). Although random signatures were not enriched across the t-SNE, 57 MSigDB signatures correlated (Pearson r > 0.75) with -and superimposed to-the above ‘B cell’ signature. These signatures comprised relevant gene sets such as ‘B cell activation’, ‘CD22-mediated BCR regulation’, ‘GO-Immunoglobulin complex’, ‘*de novo* protein folding’ and ‘somatic mutation’, fully consistent with B cell biology (**Supplementary Table 1, Figure 1**). In contrast, no signature defining non-B cell lineages were enriched in this cluster (not shown). Visualization of metabolic pathway signatures in this PBMC data set showed that myeloid cells display higher scores than lymphocytes for glycolysis, oxidative phosphorylation (OXPHOS) and tricarboxylic acid (TCA) cycle but not fatty acid synthesis (**Supplementary Figure 1**).

**Figure 1.**
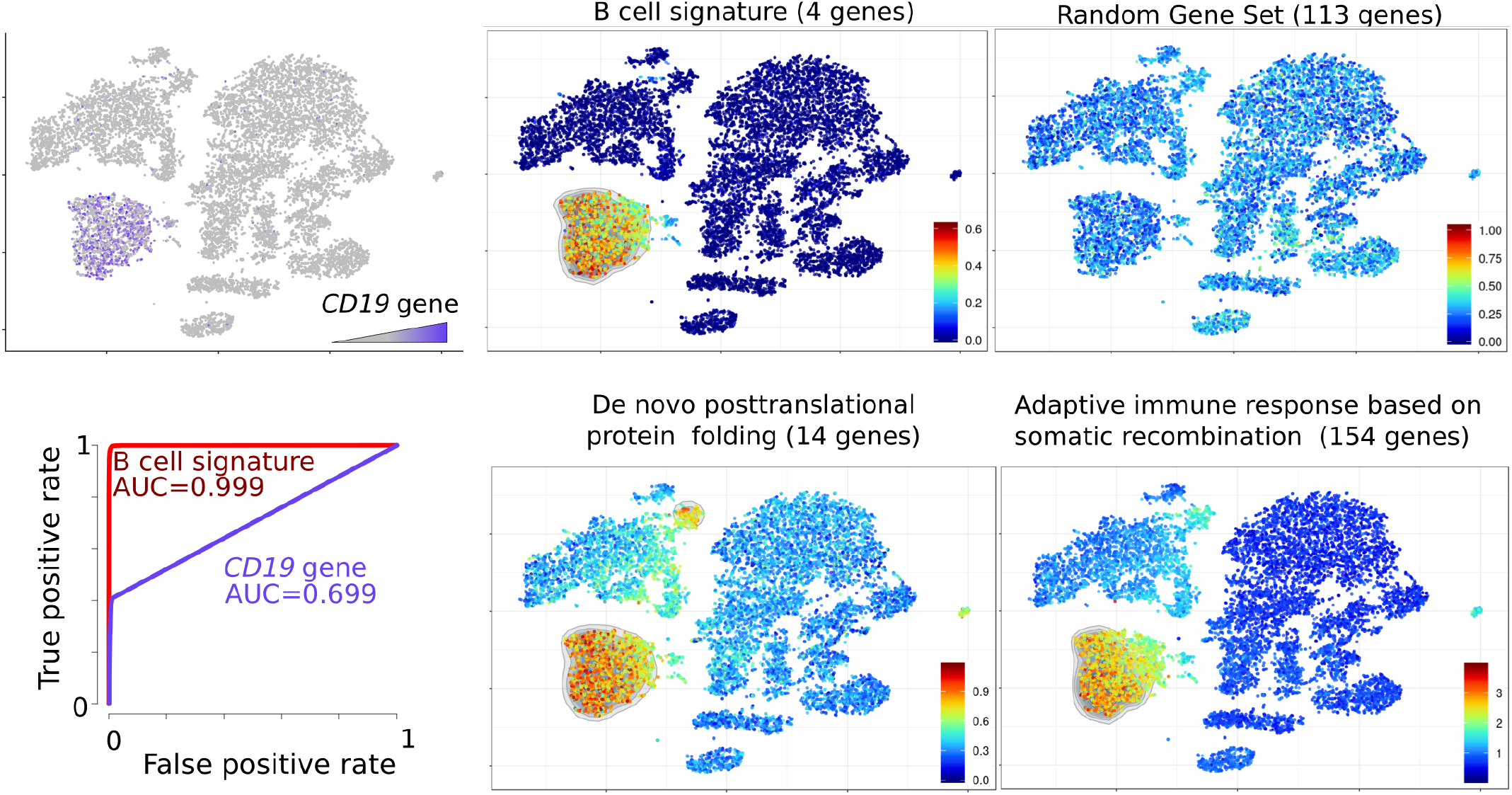
Visualization of signatures in the t-SNE map of 12k PBMC from an healthy individual. *Top left*: expression of the CD19 gene. *Top and bottom right*: scores for “B cell signature”, “random gene set”, “*de novo* post-translational protein folding” and “adaptive immune response based on somatic recombination of immune receptors built from immunoglobulin superfamily domains” (both from Gene Ontology) displayed on the t-SNE map. ROC curve (*Bottom left*) of B cell identification performance by using either the *CD19* gene alone (violet curve) or the B cell signature score (red curve).

Since the intra-tumoral microenvironment affects the metabolic reprogramming of immune cells (reviewed in (18)), such biases were explored in a cancer data set comprising matched samples of malignant and adjacent non-malignant tissues. The scRNA-seq data sets of 19k cells from 20 tumor and 4 matched normal adjacent biopsies from 6 lung cancer patients(4) were downloaded from Array Express E-MTAB-6149 and E-MTAB-6653, processed, integrated and their t-SNE was plotted. Each of these 19k single cells was also scored for the MSigDB gene sets as above. Visualizing scores for a series of lineage-defining gene sets identified the cell types of each cluster (**Supplementary Figure 2**) consistent with single gene expressions (4). The myeloid cells from either the tumor or the adjacent tissue were further gated in parallel analyses, and their scores for the above metabolic signatures were visualized. This evidenced the higher glycolytic signature of a fraction of intra-tumoral myeloid cells relative to those from adjacent lung tissue, while all cells scored quite similarly for the other pathways. Furthermore, the lung cancer cells also displayed this glycolytic bias (**Supplementary Figure 3**). Hence, scoring any single signature across the t-SNE of a scRNA-Seq data set allows to visualize its imprint at the single cell level, as exemplified above with cell identities or biological hallmarks.

Further, this strategy enabled the exploration of two signatures combined through simple subtraction or addition of their scores, and projecting the result on the t-SNE map. This was illustrated using the human testis cell atlas scRNASeq data set of 6.5 k cells involved in adult spermatogenesis (5). This publicly available data set was downloaded, processed as above and its partitioning into 16 clusters was projected onto a t-SNE of the germ cells pseudo-time maturation trajectory. Both cluster identities (5) and positions on the trajectory (**Figure 2a**) were consistent with the molecular transitions underlying this development. The “embryonic organ development” signature was observed in spermatogonial stem cells (SSC) and their cell partners (testicular macrophages, Leydig, Sertoli, myoid and endothelial cells), while “male meiosis” and “spermatogenesis” signatures hit only the early primary spermatocytes and mature sperm cells, respectively. The “GO_mitochondrion” signature gradually decreased along the spermatocyte maturation, contrasting with the other microenvironmental cells (**Supplementary Figure 4a**). Whether the spermatogenesis energy is fueled by mitochondrial oxidative phosphorylation, or by glycolysis, or both continuously, or even by their stage-specific complementation, is matter of debates. Here, displaying across the same data set the combined score for “GO_Glycolysis” gene set *minus* “GO_OXPHOS” gene set illustrated the glycolytic imbalance which progressively settles along the sperm cell maturation (**Figure 2b**). This was consistent with the above-depicted decrease of mitochondria in maturing spermatozoids (**Supplementary Figure 4a**) and the progressive catabolic switch of glucose metabolism evidenced by the combined score for “GO_Glucose metabolic process” gene set *minus* “GO_Glucose catabolic process” gene set (**Supplementary Figure 4b**). Thus Single-Cell Signature Explorer allows to visualize combinations of 2 distinct signatures, yielding newer informative displays, possibly completing those from single gene set-based signatures.

**Figure 2.**
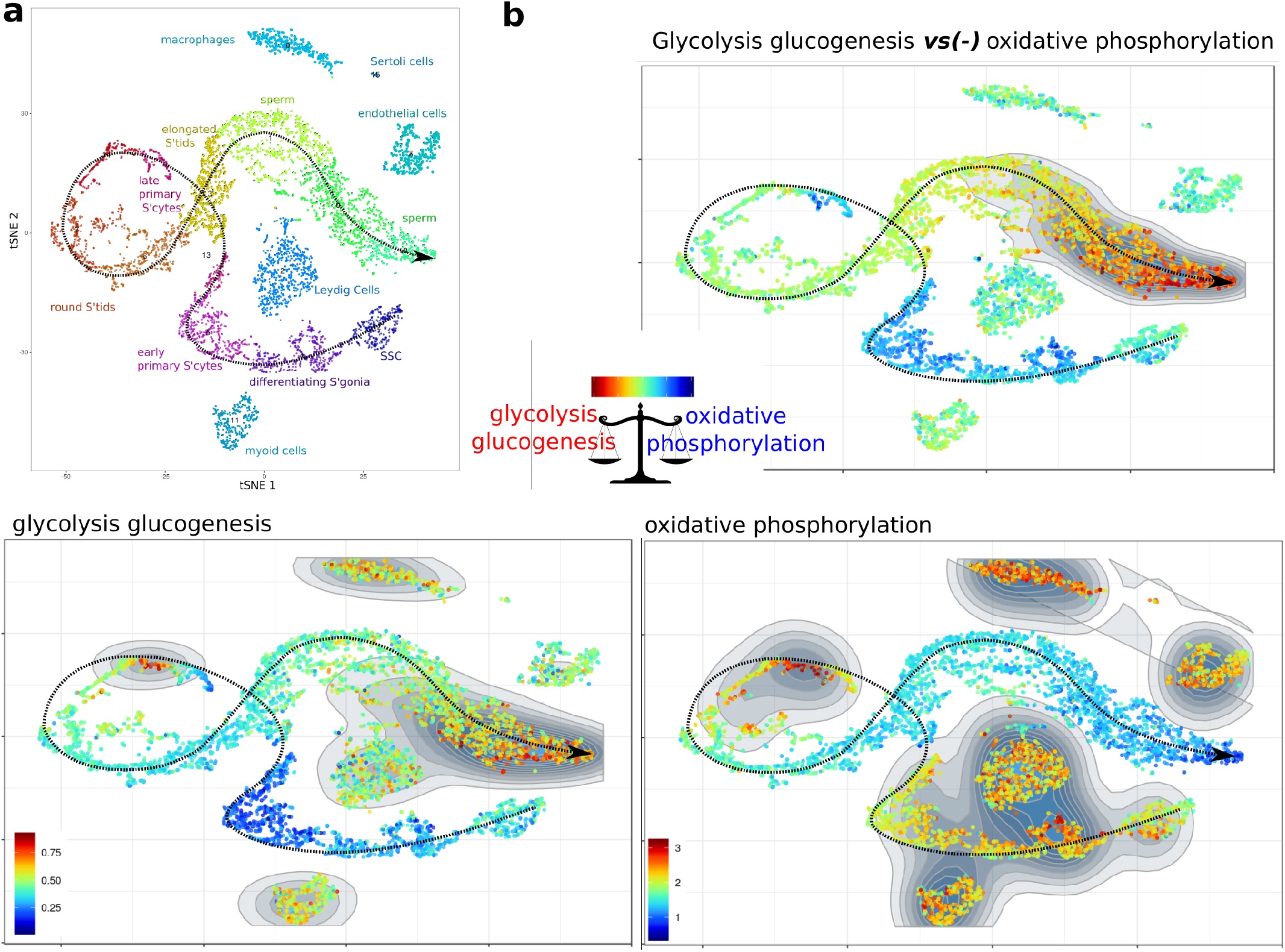
Single and combined signatures of metabolic pathways in human adult testis cells. SSC: spermatogonial stem cells. **a**: Human adult testis cell types and their maturation trajectory t-SNE. **b**: (*bottom panels*) Single signatures of “glycolysis glucogenesis” (KEGG, 62 genes), and “oxidative phosphorylation” (KEGG, 136 genes), and (*top panel*) their combined signature using the “*minus*” operator on the t-SNE map.

The recently published algorithms Seurat’s Cell CycleScore module (7), and AUCell (14) can also compute the enrichment scores of gene set-based signatures from single cell transcriptomes. With the above-depicted healthy PBMC, lung adenocarcinoma tumors, and human adult testis data sets, we compared Single-Cell Signature Explorer to these latter and to GSVA/ssGSEA on criteria including high throughput performance, computing time, and display of the results (**Supplementary Table 2 and Supplementary Figure 5**). While Seurat CellCycleScoring function computes almost as quickly, and yields similar results, as Single-Cell Signature Explorer (**Supplementary Figure 6**), it only scores few gene sets pairs with anti-correlated expression, and does not display t-SNE plots of the results. GSVA/ssGSEA, incepted for scoring gene sets from bulk transcriptomes but not zero-inflated single cell transcriptomes, requires such tedious computing times for a single signature that it cannot perform massively parallel scoring of large collections of gene sets, and does not display t-SNE results. Although the gene rank-based AUCell algorithm computes scores 5-times faster than GSVA/ssGSEA, it remains 30-times slower than Signature Explorer, and does not display interactive t-SNE maps of the resulting scores. In addition to its computing performance, the versatility of Single-Cell Signature Explorer permits analysis of data from various scRNA-Seq and sequencing platforms (**Supplementary Figure 7**). Notably, by uploading the gene set databases for macaccus, mouse, rat, and zebra fish (https://sites.google.com/site/fredsoftwares/products/databases) into Single-Cell Signature Scorer, it allows exploration of non-human samples. Thus Single-Cell Signature Explorer represents the reference tool for general exploration of scRNASeq data sets.

## CONCLUSION

Functionally meaningful displays of gene set-based signature enrichment are essential to understand t-SNE maps of complex cell samples, but so far, no current method perform this rapidly for a plethora of signatures at the single cell level across large scRNAseq data sets. By quickly delineating multi-gene features such as cell lineage or metabolic pathways with Single-Cell Signature Explorer in lung tumours and normal human blood and testis, we showed that gene set-based signatures outperform single genes and provide a straightforward visualization of sample’s hallmarks. Within few minutes from any computer, this new method provides users with thousands of gene set-based signatures for thousands of single cells, and the immediate visualization in t-SNE of any of these single signatures or combination of signatures. Being compatible with any cell sample, scRNASeq platform and sequencing tools, and any gene set either used-defined or from MSIgDB’s current 17,810 signatures, its applications are as broad as the scRNASeq technology itself.

## ACKNOWLEDGEMENTS

Aviv Regev and Asaf Madi (Broad Institute) are acknowledged for kindly sharing R scripts for density plots, and Eric Espinosa (CRCT) for providing the mastocyte gene set.

