## Supplementary Information for "Single-Cell Signature Explorer for comprehensive visualization of single cell signatures across scRNA-seq data sets"

### Supplementary methods

### scRNAseq DataSet Noise Reduction

Single-Cell Signature Explorer indifferently processes scRNAseq datasets composed of either raw UMI data, noise-reduced transformed UMI data, or UMI data normalized by Seurat or by other pipelines. User must select accordingly the type of data set to be further analysed by Single-Cell Signature Explorer. Single cell transcriptomic technologies produce scRNAseq datasets in which technical noise may hinder biological variability. This technical noise may result from both gene sampling fluctuations and cell-to-cell variations in sequencing efficiency[1].

Variations in gene sequencing efficiency can be reduced by median normalization of the total UMI counts per cell for all cells of the data sets, and re-scaling the expression profile of each gene from each cell as follows :

$UMI_{MN} = UMI \times \frac{t_{Med}}{t_{Cell}}$

where $t_{Cell}$ is the total UMI sum for a cell, $t_{Med}$ is the data set's median of all $t_{Cell}$, and $UMI_{MN}$ is the UMI corrected by median normalisation.

Second, gene sampling noise, which obeys Poissonian statistics[1], may be stabilized by the Freeman-Tuckey transform[2] of the above median-normalized UMI counts :

$UMI_{FTMN} = \sqrt{UMI_{MN}} + \sqrt{UMI_{MN} + 1}$

where $UMI_{FTMN}$ is the UMI corrected by both median normalization and Freeman-Tuckey transform. We provide below a comparative analysis by Single-Cell Signature Explorer of the same 8k PBMC data set under the form of either raw UMI data, Seurat-normalized data, or Noise-Reduced (FTMN) data computed by scRNAseq DataSet Noise Reductor.

**1-Data noise reduction**

The total UMI counts of each single cells from 8k PBMC data set (downloaded from 10xGenomics website) were compared when using either the raw UMI, the FTMN noise reduced UMI, or the Seurat-normalized UMI. As expected, the noise reduction with FTMN reduced and smoothed the total per cell across the data set, yielding a CV of 18.3% instead of 37.9% with raw data, and 13.9 % with Seurat normalization (Suppl. Figure 8). Then, a single cell was randomly selected from the same data set and its transcriptomes based on either raw UMI , FTMN (noise reduced) UMI or Seurat-normalized UMI data from were compared. This evidenced a clear-cut and global reduction of the FTMN-transformed UMI data relative to the raw UMI, consistent with the previous conclusion (Suppl. Figure 8). Hence FTMN effectively reduced the CV of the raw UMI data at the price of their global reduction.

**2-Impact of data noise reduction on signature t-SNE maps**

We then compared the analysis by Single-Cell Signature Explorer of the same 8k PBMC data set (downloaded from 10xGenomics website) when computed from the raw UMI data, FTMN (noise reduced) UMI data or Seurat-normalized UMI data. With the 8k PBMC, the computing time for scoring all the 186 gene sets from the KEGG database was of 23 sec. for the raw UMI data, 22 sec. for the Seurat-normalized UMI data and 524 sec. (8 min 24sec) for the FTMN-transformed UMI data (Supplementary table 6).

These scorings yielded similar t-SNE images of signatures for 3 representative KEGG metabolic pathways, respectively composed of highly expressed genes (OXPHOS, 135 genes), medium-expressed genes (Glycolysis, 62 genes) or weakly-expressed genes (Arginine and proline metabolism, 54 genes). Both Pearson and Spearman correlation coefficients between the 8k scores from each pair of data set type are provided for each metabolic pathway. All pairs were strongly correlated. The lowest correlation (Pearson r= 0.88) was observed between the Raw UMI and Noise-Reduced UMI data sets for the largest gene set OXPHOS, mostly owing to their different ratings of OXPHOS in the small cell clusters at the uppest, lowest and left sectors of the t-SNE (Supplementary figure 9). No such discrepancy was noticed for the 2 others metabolic pathways, despite consistently lower levels of expression across the t-SNE.

The same comparative study was then focused to cell lineage-defining signatures for 3 abundant, well-defined, and different cell types: B cells (4 genes), NK cells (48 genes) and macrophage cells (36 genes). Both t-SNE images of these signatures and correlation coefficients of all pairs of data sets were strongly consistent in demonstrating that Single-Cell Signature Explorer yields similar results from these 3 types of data.

Finally the same comparative study was done for 3 random signatures composed of randomly chosen sets of genes of various sizes. As above, both t-SNE images of these signatures and correlation coefficients of all pairs of data sets were strongly consistent in yielding the same results whatever the type of data used. The lowest correlation observed (Pearson r= 0.86) was again observed between the Raw and the Noise-Reduced datasets for the largest (462 genes) random data set, while the smallest random gene set yielded very strongly correlated signatures across the t-SNE (Spearman r= 1).

We then investigated the impact of data noise reduction on the signature scores. For a given signature defining positive and negative cells, here B cell (Supplementary figure 10), we then compared the distribution patterns of single cell scores across different clusters of the t-SNE. As expected for B cells *vs* non-B cell clusters, this signature displayed clearly different score means across the t-SNE clusters. This comparison evidenced consistently lower scores (*i.e.* 3-10 times lower means and ranges) with FTMN data than with raw or Seurat normalized data. On the other hand however, with noise reduced data the contrast of signature scores in each cluster decreased and their homogeneity increased. The same type of analysis was done for a metabolic signature such as glycolysis. Although in this case, the cluster means differed less between clusters (Supplementary figure 11), the means and ranges of all the clusters scores were also reduced with noise reduced data. As previously, the contrast of scores in each cluster decreased and their homogeneity increased with the noise-reduced UMI data.Thus, whatever the signature, Single-Cell Signature Explorer produces similar t-SNE images from either raw, noise-reduced, or Seurat normalized data.

Altogether, we concluded that Single-Cell Signature Explorer does not necessarily require noise-reduction from the scRNASeq data sets, and produces similar t-SNE images from raw UMI, noise-scores reduced UMI, or Seurat normalized UMI data, at the price of considerably longer computing time for the noise-reduced data.

### Supplementary figures

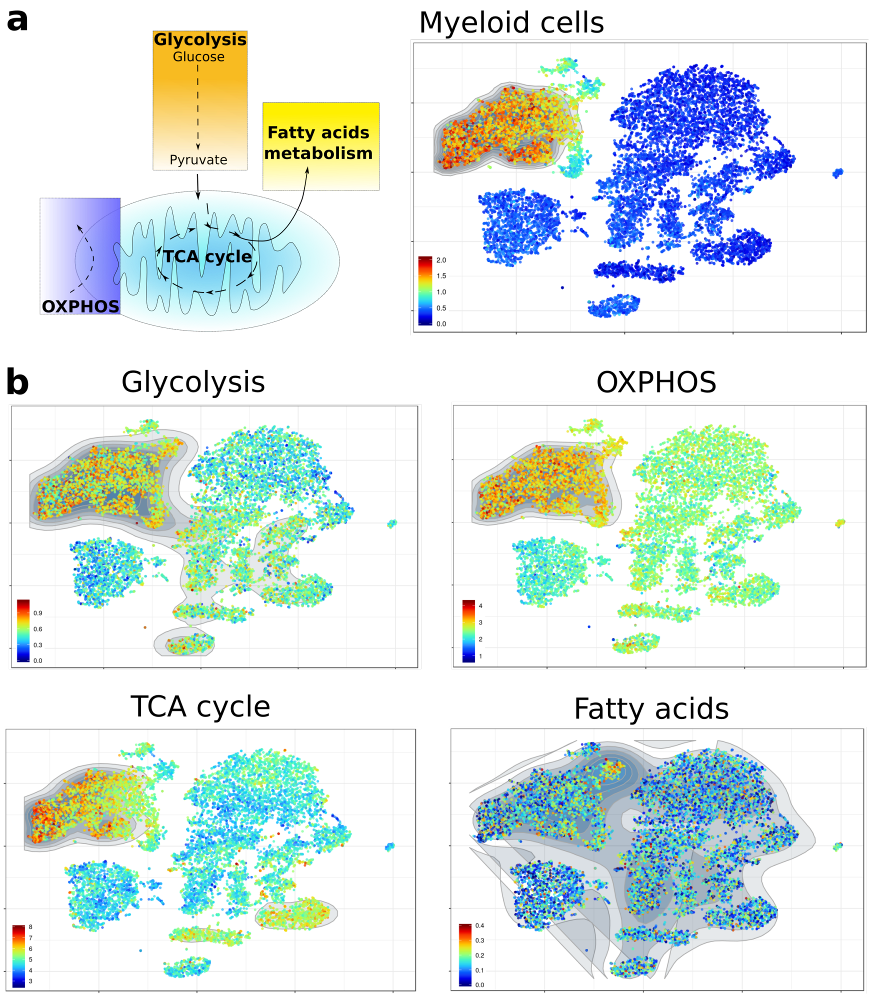


**Supplementary Figure 1.** Visualization of single cell PBMC scores for metabolic pathways unveils the higher signatures of myeloid cells for glycolysis (KEGG glycolysis and gluconeogenesis, 63 genes) , oxydative phosphorylation (OXPHOS) (KEGG Oxidative phosphorylation, 136 genes) and tricarboxylic acid (TCA) (KEGG citrate cycle-TCA cycle, 32 genes), but not for fatty acid metabolism (KEGG fatty acid metabolism, 42 genes.)

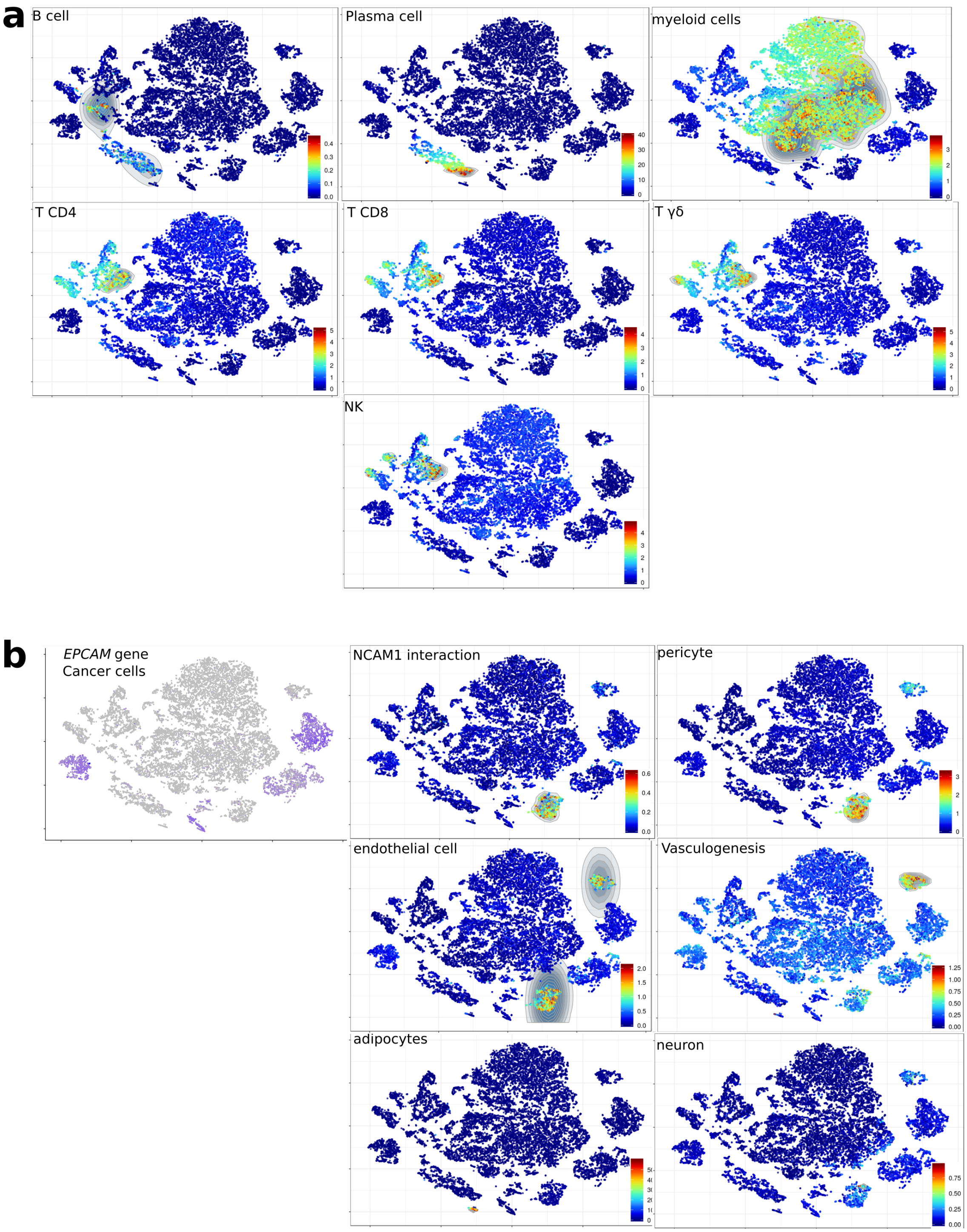


**Supplementary Figure 2.** Identification of cell types in the t-SNE of 19k cells from lung adenocarcinoma tumors and normal adjacent tissue (data from Array Express E-MAAB-6149 and E-MTAB-6653[3]). "B cell" (4 genes), "plasma cell" (47 genes), myeloid cells (31 genes), T CD4 (47 genes), T CD8 (38 genes), T $\gamma\delta$ (52 genes), Natural Killer (NK, 48 genes), "NCAM1 interaction" (reactome, 36 genes), pericyte (47 genes), endothelial cells ( 34 genes), "vasculogenesis" (Gene Ontology, 59 genes), adipocyte (48 genes), neuron (49 genes) pathways were scored and mapped. These gene sets are detailed in **Supplementary Table 5**.

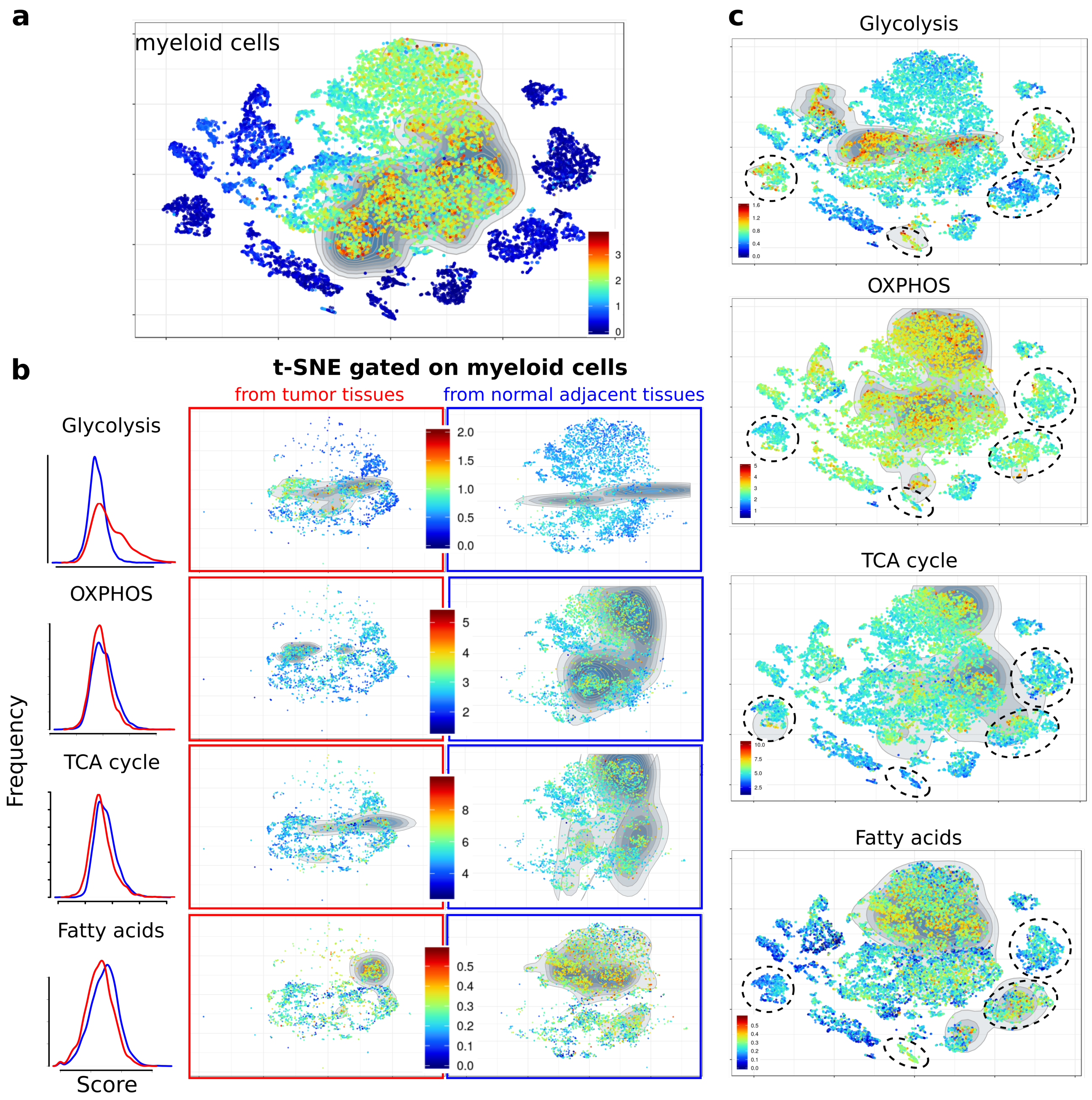


**Supplementary Figure 3.** Metabolic bias of myeloid cells in lung cancer. a: Myeloid cells in the lung adenocarcinoma tumors and normal adjacent tissue data set shown in Suppl Fig. 2. b:Single cell scores for the specified metabolic pathways in myeloid cells (gated shown in panel a) from either the tumor (middle panels) or adjacent normal tissue (right panels). Distribution of single myeloid cell scores for the corresponding pathways (left panel). c: Same pathways visualized across the entire t-SNE featuring cancer cell clusters (dotted circles).

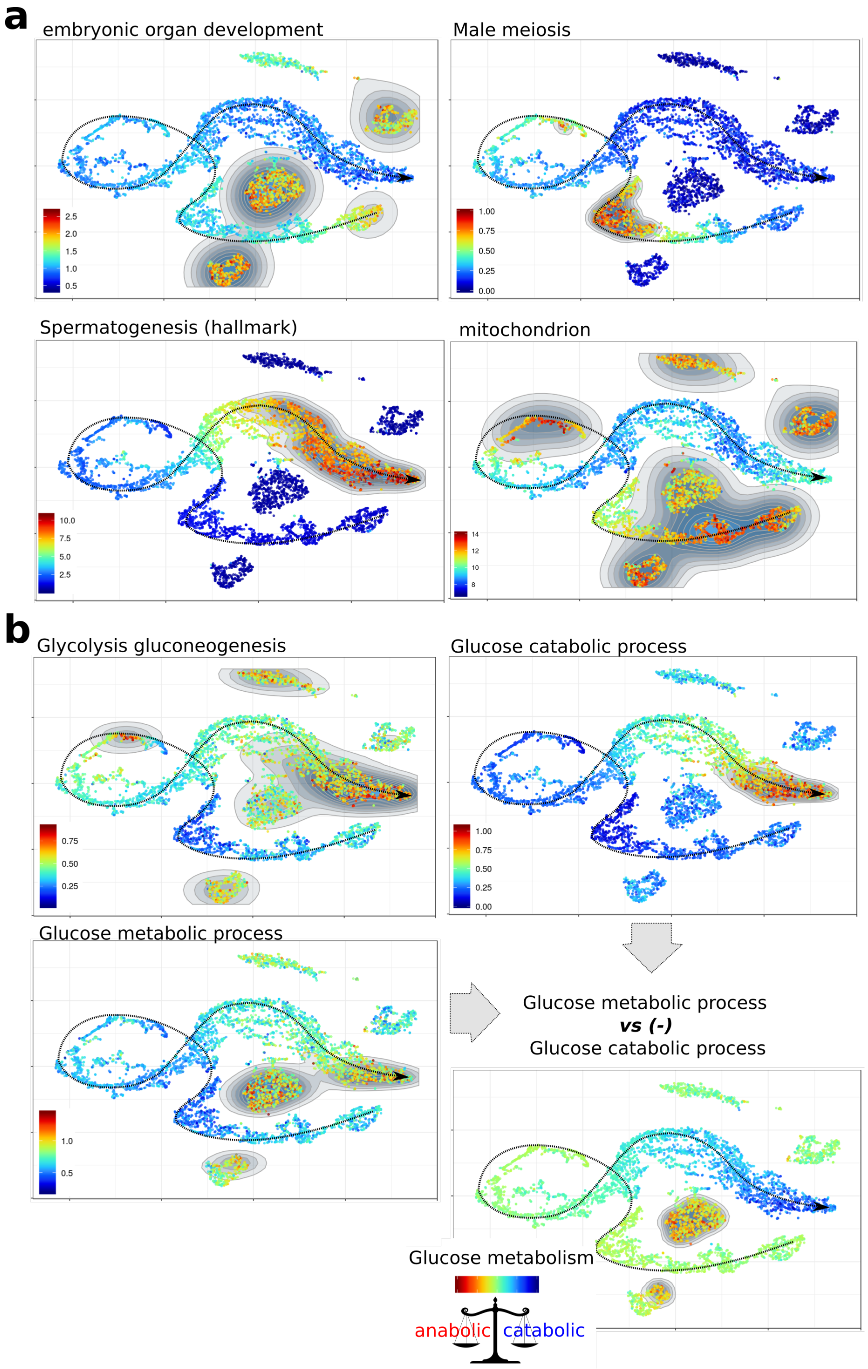


**Supplementary Figure 4.** Visualization of signatures in the t-SNE map of 6.5k cells involved in adult spermatogenesis. Scores for "embryonic organ development" (Gene Ontology, 406 genes), "male meiosis" (Gene Ontology, 39 genes), "spermatogenesis" (Hallmark, 135 genes), "mitochondrion" (Gene Ontology, 1633 genes), "glycolysis glycogenesis" (KEGG, 62 genes), "glucose catabolic process" (Gene Ontology, 29 genes) and "glucose metabolique process"(Gene Ontology 119 genes) or their combination were displayed on the t-SNE map.

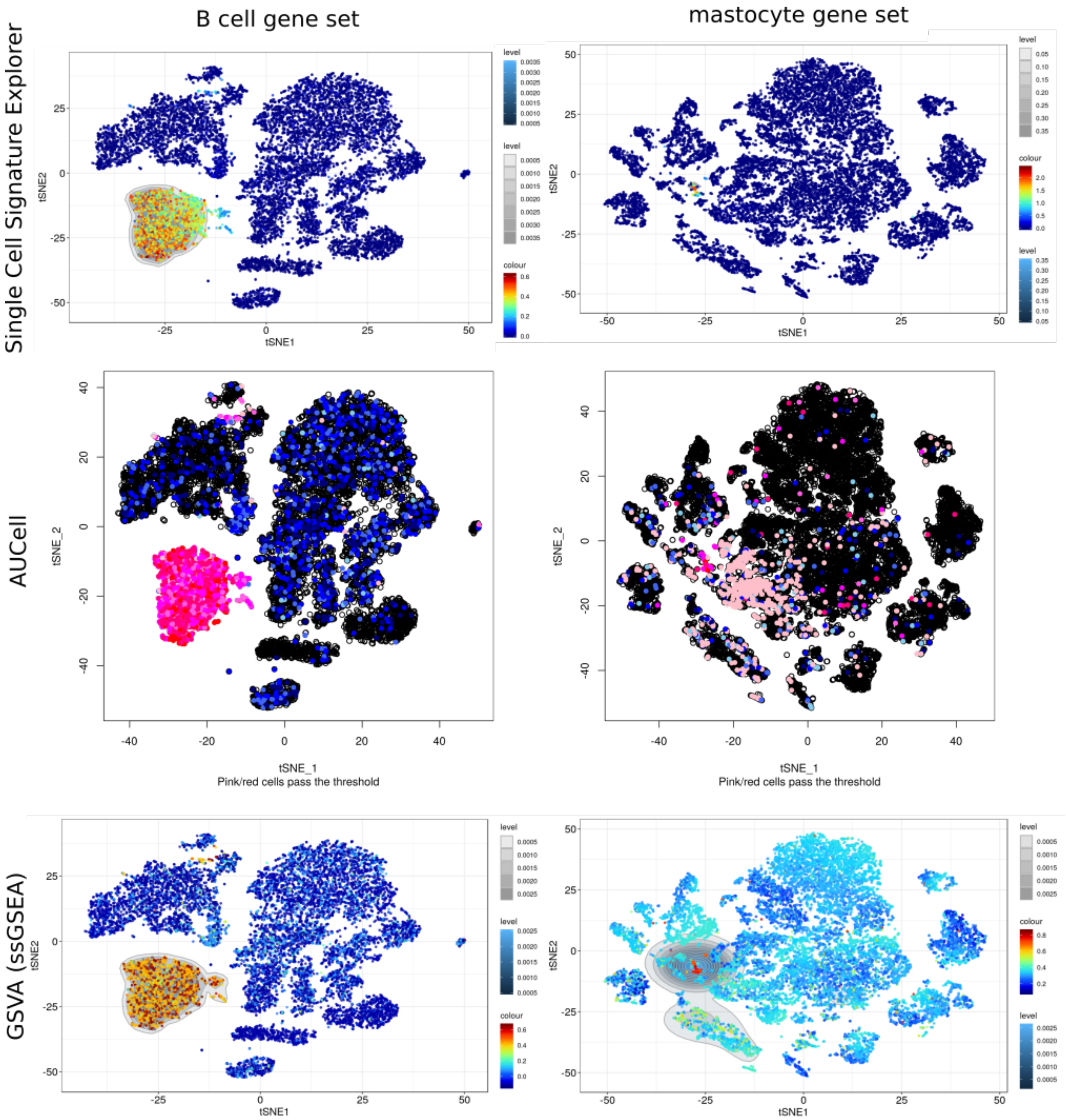


**Supplementary Figure 5.** Visualization of signatures on a t-SNE map of 12k PBMC from an healthy individual *(left)* and 19k cells from lung adenocarcinoma tumors and normal adjacent tissue (*right*). *Top*: scores calculated with Single-Cell Signature Explorer *middle*: scores calculated with AUCell *bottom*: scores calculated with GSVA/ssGSEA. Note that both results from GSVA/ssGSEA could only be visualized using Single-Cell Signature Viewer.

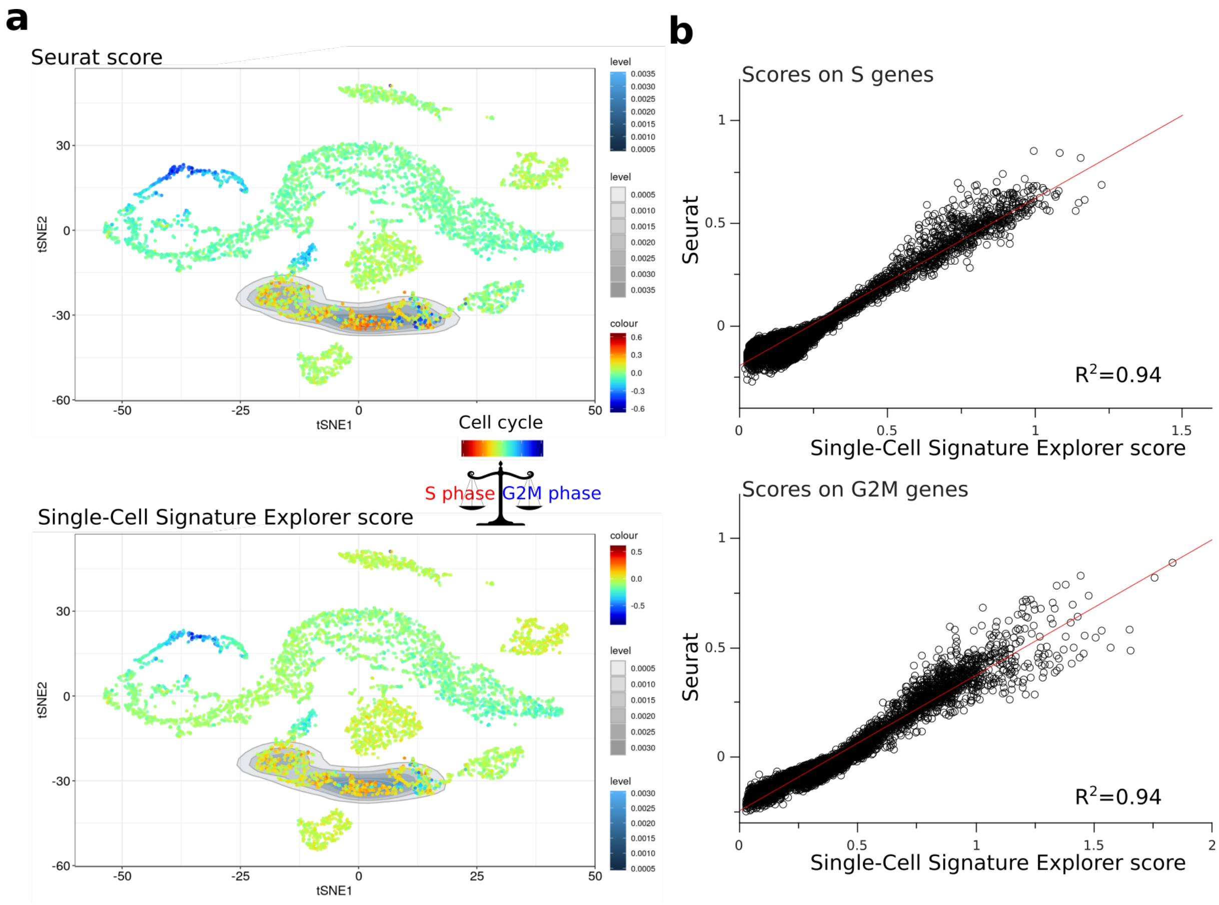


**Supplementary Figure 6.** Visualization of the cell cycle phases S and G2M (with signatures as defined in the Seurat manual) on the t-SNE map of human adult spermatogenesis (6.5k cells). *a*: Scores calculated using the Seurat CellCycleScore package *(Top)* or Single-cell Signature Explorer *(Bottom)*. The visualization of Seurat CellCycleScore results was done by using Single-cell Signature Explorer. *b*: Correlation of scores calculated by Seurat or Single-cell Signature Explorer across the data set.

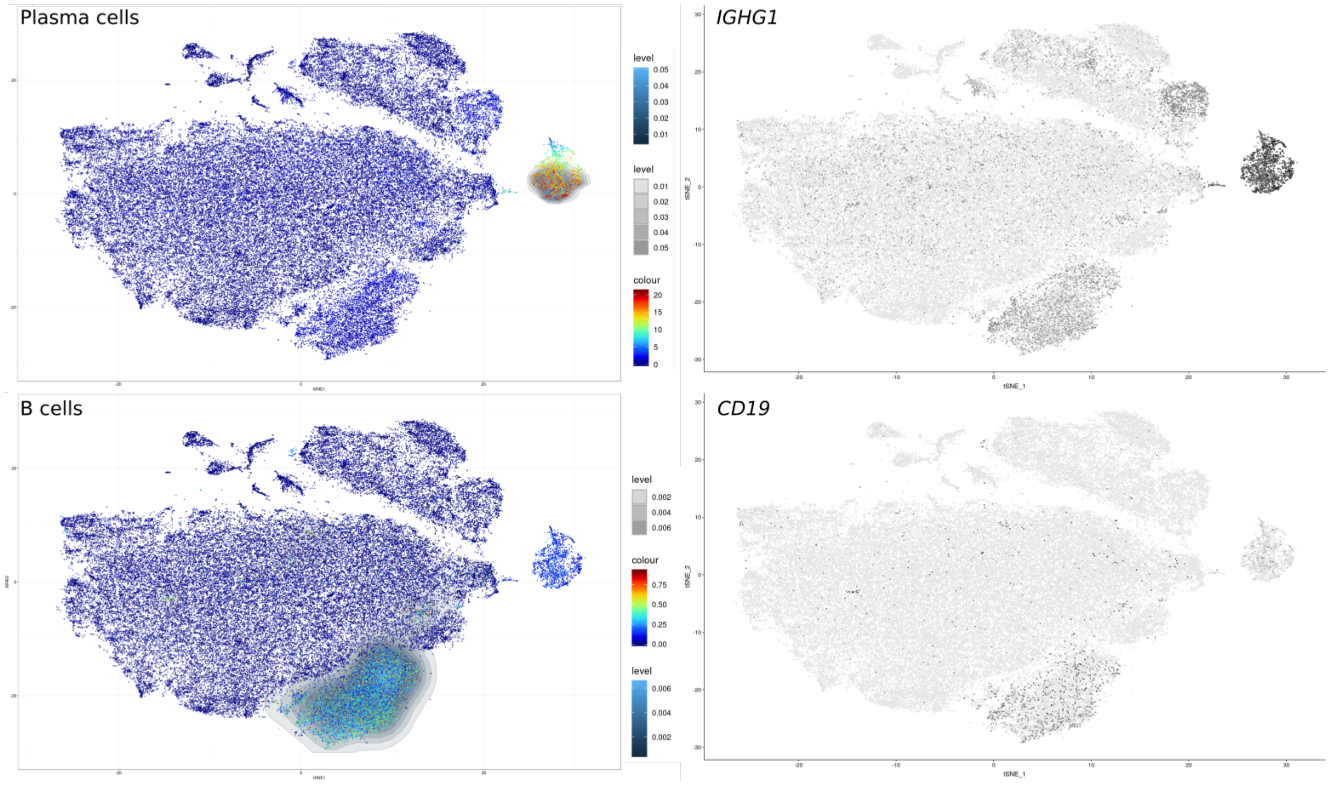


**Supplementary Figure 7.** Visualization of B cells and plasma cells signatures on the t-SNE map of 64k cells in melanoma (GSE123139) produced by the MARS-seq technology[4]. *left*: Scores calculated by Single-Cell Signature Scorer *right*: Single genes defining these two populations as described in[4]

.

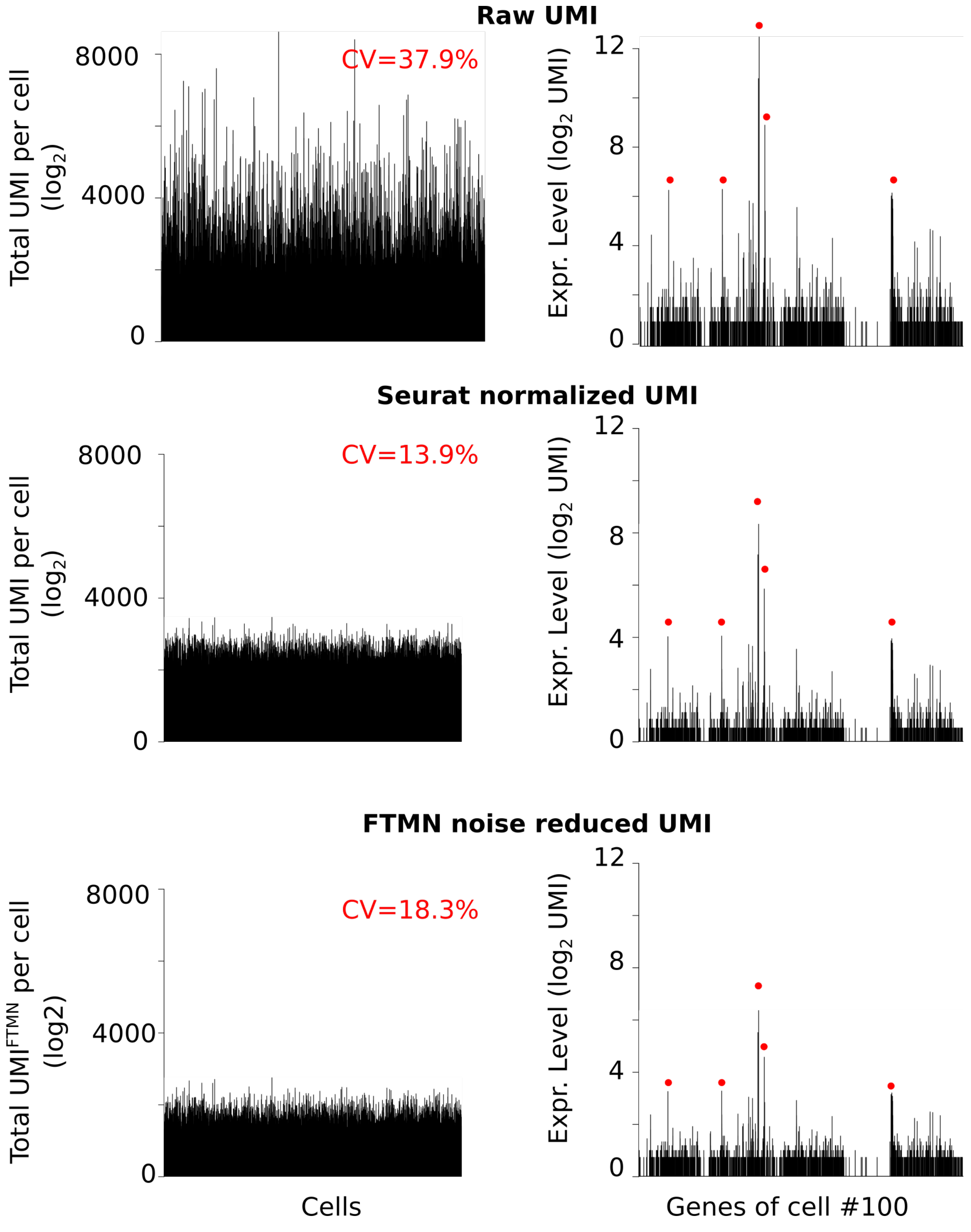


**Supplementary Figure 8.** Transformation of raw UMI data by Seurat normalization or by noise reduction with FTMN transform data for noise reduction. Left: Total of gene expression across the cells. Coefficients of variation (CV) are indicated in red. Right: Profile of one cell (#100) across gene expression. Red dots indicated the same 5 prominent genes in the 3 data set

Raw FTMN Seurat

**Oxidative phosphorylation**

| | | R Pearson | R Spearman |
|---|---|---|---|
| raw | FTMN | 0.8819 | 0.8999 |
| raw | Seurat | 0.9669 | 0.9623 |
| FTMN | Seurat | 0.8938 | 0.9161 |

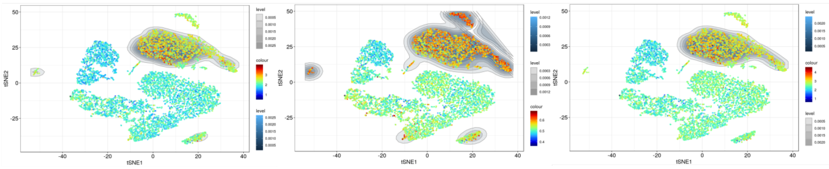

**Glycolysis gluconeogenesis**

| | | R Pearson | R Spearman |
|---|---|---|---|
| raw | FTMN | 0.9660 | 0.9683 |
| raw | Seurat | 0.9700 | 0.9788 |
| FTMN | Seurat | 0.9705 | 0.9789 |

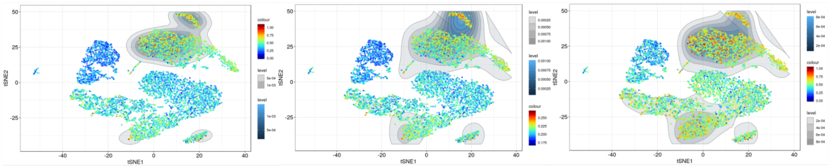

**Arginine and proline metabolism**

| | | R Pearson | R Spearman |
|---|---|---|---|
| raw | FTMN | 0.9554 | 0.9849 |
| raw | Seurat | 0.9646 | 0.9873 |
| FTMN | Seurat | 0.9922 | 0.9972 |

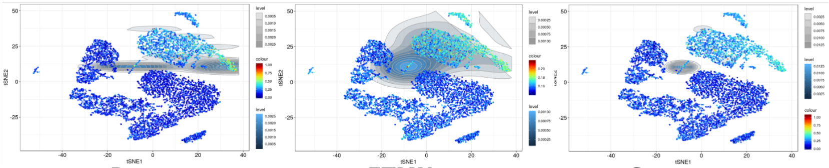

Raw FTMN Seurat

**B cells**

| | | R Pearson | R Spearman |
|---|---|---|---|
| raw | FTMN | 0.9929 | 0.9991 |
| raw | Seurat | 0.9960 | 0.9996 |
| FTMN | Seurat | 0.9958 | 0.9990 |

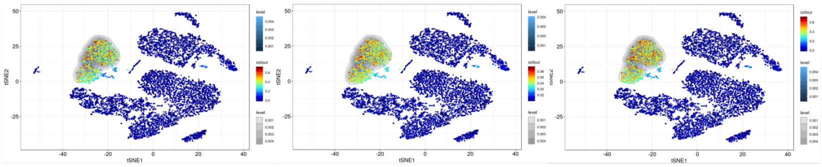

**NK cells**

| | | R Pearson | R Spearman |
|---|---|---|---|
| raw | FTMN | 0.9869 | 0.9519 |
| raw | Seurat | 0.9898 | 0.9564 |
| FTMN | Seurat | 0.9903 | 0.9671 |

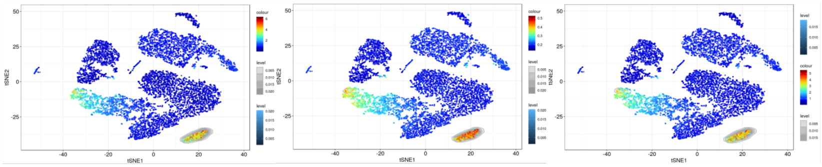

**macrophages**

| | | R Pearson | R Spearman |
|---|---|---|---|
| raw | FTMN | 0.9785 | 0.9766 |
| raw | Seurat | 0.9821 | 0.9837 |
| FTMN | Seurat | 0.983 | 0.9871 |

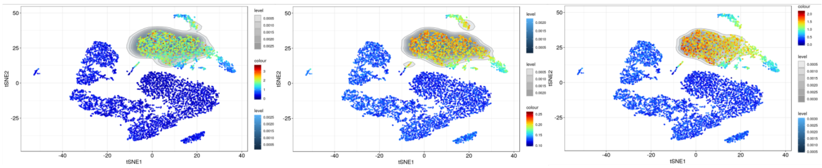

Raw FTMN Seurat

**Random Gene Set #988 (462 genes)**

| | | R Pearson | R Spearman |
|---|---|---|---|
| raw | FTMN | 0.8781 | 0.8652 |
| raw | Seurat | 0.9422 | 0.9336 |
| FTMN | Seurat | 0.9779 | 0.9793 |

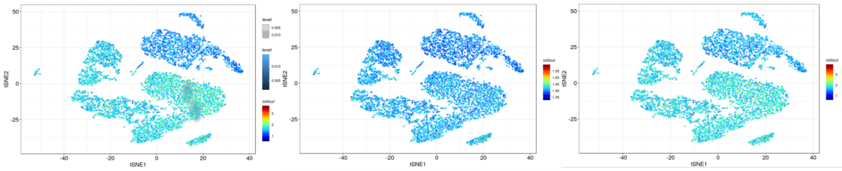

**Random Gene Set #728 (142 genes)**

| | | R Pearson | R Spearman |
|---|---|---|---|
| raw | FTMN | 0.9519 | 0.9550 |
| raw | Seurat | 0.9652 | 0.9643 |
| FTMN | Seurat | 0.9927 | 0.9954 |

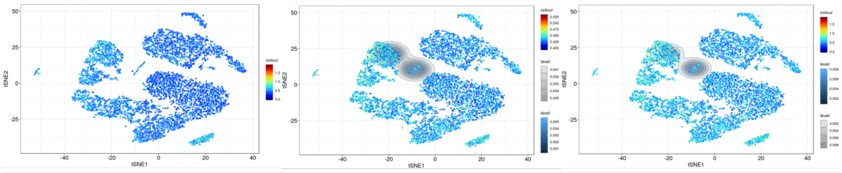

**Random Gene Set #685 (4 genes)**

| | | R Pearson | R Spearman |
|---|---|---|---|
| raw | FTMN | 0.9920 | 0.9995 |
| raw | Seurat | 0.9958 | 1.0000 |
| FTMN | Seurat | 0.9949 | 0.9995 |

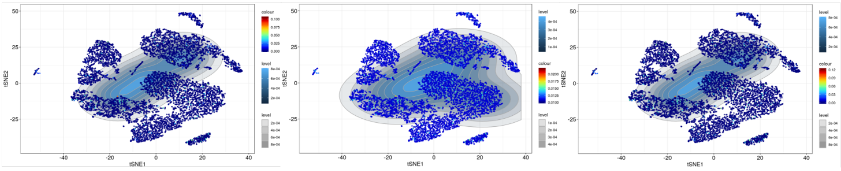

**Supplementary Figure 9.** Visualization of metabolism pathways, population signature and random gene set on the t-SNE map of 8k PBMC from healthy donor using either raw UMI counts, noise-reduced UMI counts by FTMN transform or UMI normalized by Seurat. Left: r coefficients for pearson and spearman methods between each normalization

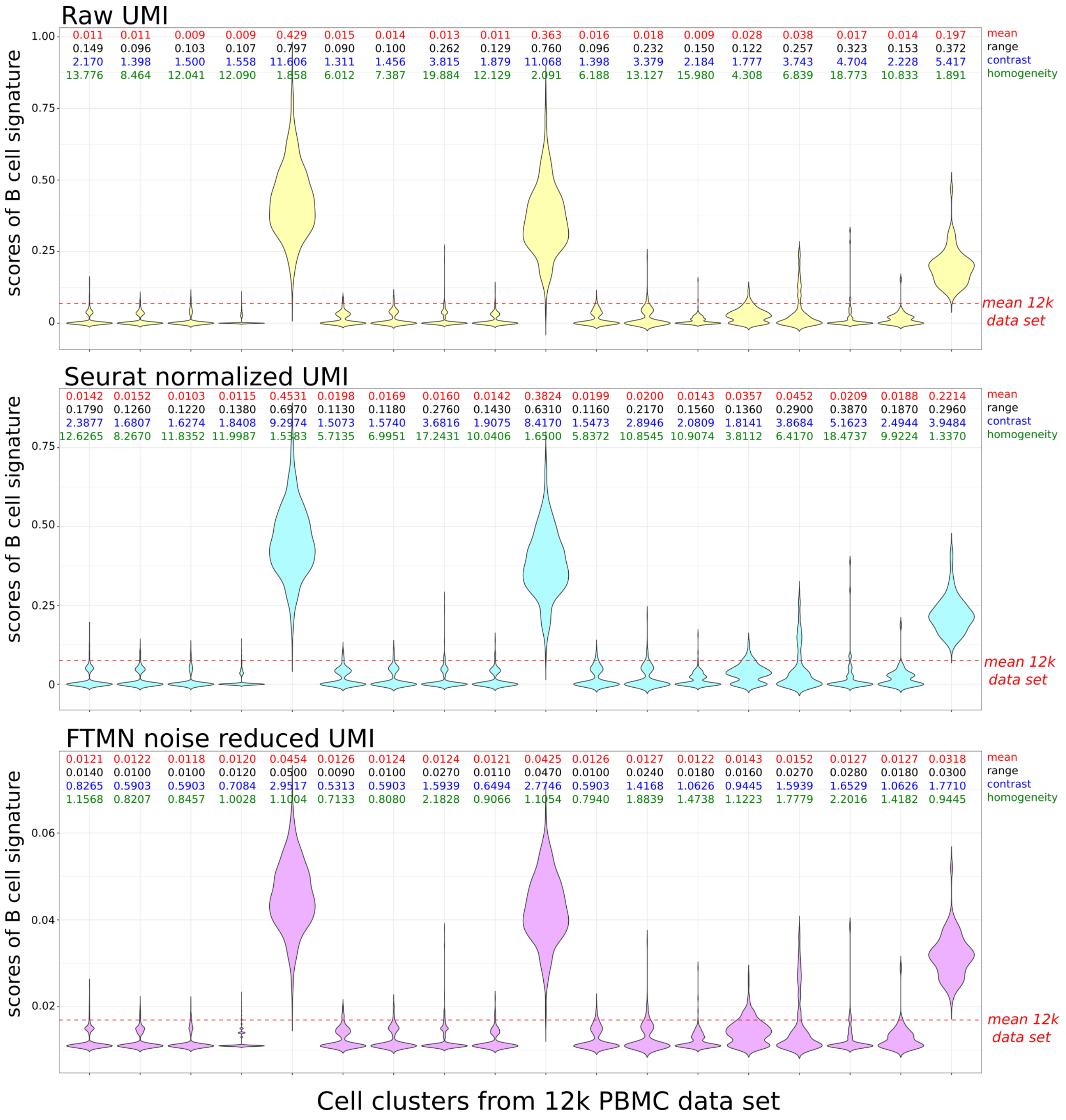


**Supplementary Figure 10.** Scores of B cell signature gene sets by cluster of 8k PBMC from healthy donor using either raw UMI counts, noise-reduced UMI counts by FTMN transform or UMI normalized by Seurat. For each data set and each cluster, the mean of the cluster, range (maximum of the cluster-minimum of the cluster), contrast (range of the cluster/mean of the data set) and homogeneity (range of the cluster/mean of the cluster) is indicated above each violin.

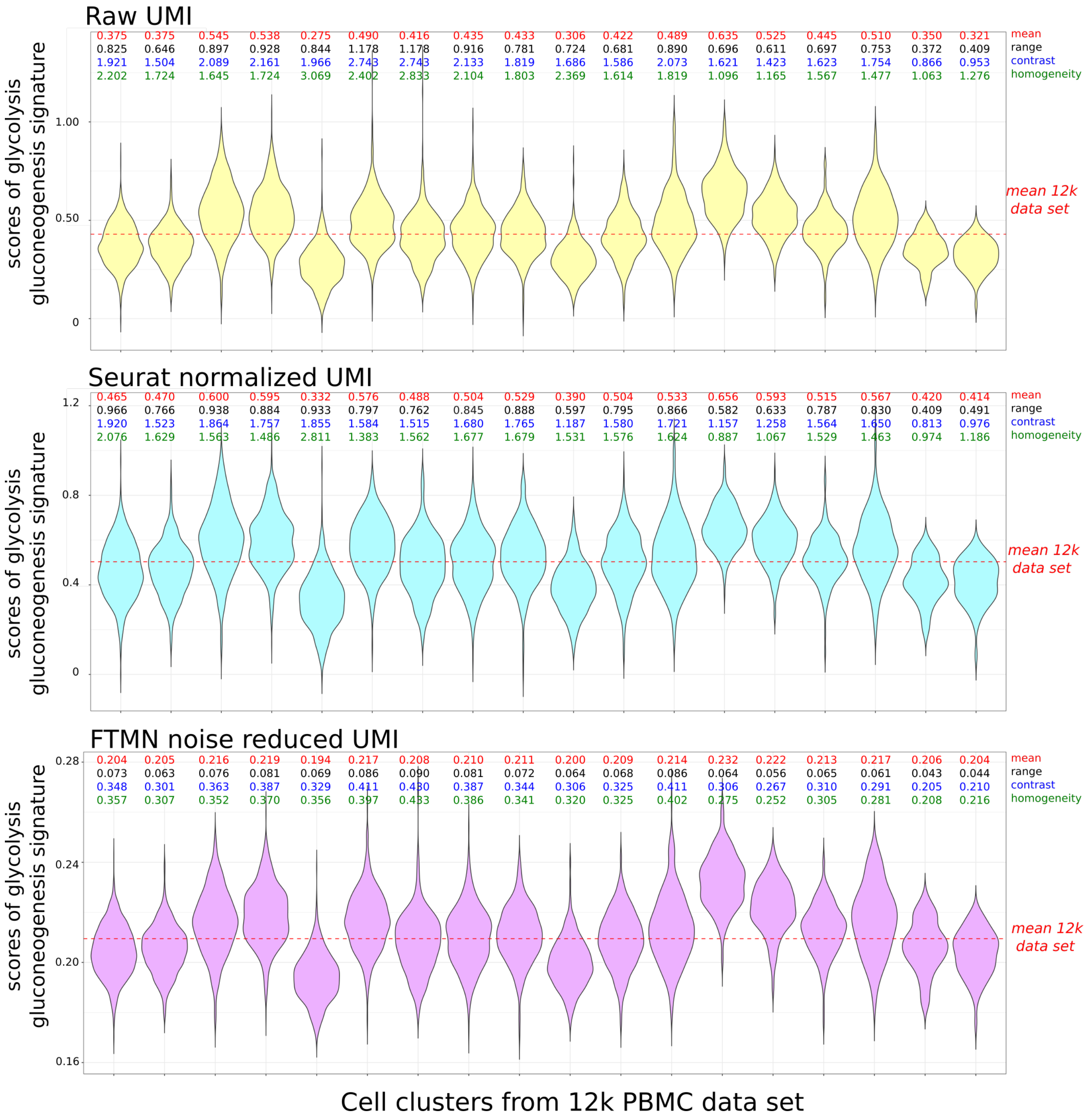


**Supplementary Figure 11.** Scores of Glycolysis gluconeogenesis gene sets (KEGG) by cluster of 8k PBMC from healthy donor using either raw UMI counts, noise-reduced UMI counts by FTMN transform or UMI normalized by Seurat. For each data set and each cluster, the mean of the cluster, range (maximum of the cluster-minimum of the cluster), contrast (range of the cluster/mean of the data set) and homogeneity (range of the cluster/mean of the cluster) is indicated above each violin.

### Supplementary Tables

| var1 | var2 | Pearson coefficient |
|---|---|---|
| B_cells | Cd22_Mediated_Bcr_Regulation | 0.9243 |
| B_cells | Go_Immunoglobulin_Complex | 0.9207 |
| B_cells | Antigen_Activates_B_Cell_Receptor_(Bcr)_Leading_To_Generation_Of_Second_Messengers | 0.9147 |
| B_cells | Go_B_Cell_Receptor_Signaling_Pathway | 0.8961 |
| B_cells | Go_Positive_Regulation_Of_B_Cell_Activation | 0.8764 |
| B_cells | Go_Humoral_Immune_Response_Mediated_By_Circulating_Immunoglobulin | 0.8724 |
| B_cells | Go_B_Cell_Mediated_Immunity | 0.8592 |
| B_cells | Classical_Antibody-Mediated_Complement_Activation | 0.8554 |
| B_cells | Regulation_Of_Complement_Cascade | 0.852 |
| B_cells | Go_Immunoglobulin_Receptor_Binding | 0.8497 |
| B_cells | Go_Regulation_Of_B_Cell_Activation | 0.8465 |
| B_cells | Go_Adaptive_Immune_Response_Based_On_Somatic_Recombination<br>_Of_Immune_Receptors_Built_From_Immunoglobulin_Superfamily_Domains | 0.8404 |
| B_cells | Role_Of_Lat2ntallab_On_Calcium_Mobilization | 0.8375 |
| B_cells | Scavenging_Of_Heme_From_Plasma | 0.8318 |
| B_cells | Go_Antigen_Binding | 0.8274 |
| B_cells | Go_Lymphocyte_Mediated_Immunity | 0.8211 |
| B_cells | Go_Phagocytosis_Recognition | 0.8188 |
| B_cells | Signaling_By_The_B_Cell_Receptor_(Bcr) | 0.8179 |
| B_cells | Go_Adaptive_Immune_Response | 0.8155 |
| B_cells | Go_Regulation_Of_Macrophage_Activation | 0.8131 |
| B_cells | Go_Positive_Regulation_Of_Type_2_Immune_Response | 0.8103 |
| B_cells | Creation_Of_C4_And_C2_Activators | 0.8102 |
| B_cells | Go_Antigen_Receptor_Mediated_Signaling_Pathway | 0.8102 |
| B_cells | Go_Negative_Regulation_Of_B_Cell_Apoptotic_Process | 0.8088 |
| B_cells | Role_Of_Phospholipids_In_Phagocytosis | 0.8062 |
| B_cells | Go_Complement_Activation | 0.8061 |
| B_cells | Go_Protein_Activation_Cascade | 0.8007 |
| B_cells | Go_Mhc_Class_Ii_Protein_Complex_Binding | 0.7975 |
| B_cells | Go_Mhc_Protein_Complex_Binding | 0.7954 |
| B_cells | Go_Humoral_Immune_Response | 0.7914 |
| B_cells | Fcgr_Activation | 0.7906 |
| B_cells | Go_External_Side_Of_Plasma_Membrane | 0.7899 |
| B_cells | Go_Protein_Binding_Involved_In_Protein_Folding | 0.7872 |
| B_cells | Go_Negative_T_Cell_Selection | 0.7859 |
| B_cells | Fceri_Mediated_Ca+2_Mobilization | 0.7858 |
| B_cells | Go_De_Novo_Posttranslational_Protein_Folding | 0.7828 |
| B_cells | Go_Regulation_Of_B_Cell_Apoptotic_Process | 0.7815 |
| B_cells | Go_Positive_T_Cell_Selection | 0.7788 |
| B_cells | Go_Negative_Regulation_Of_Dna_Damage_Response_Signal_Transduction_By_P53_Class_Mediator | 0.7779 |
| B_cells | Go_Positive_Regulation_Of_Antigen_Processing_And_Presentation | 0.7771 |
| B_cells | Go_Multivesicular_Body | 0.7765 |
| B_cells | Go_Prostanoid_Biosynthetic_Process | 0.7756 |
| B_cells | Go_Leukocyte_Mediated_Immunity | 0.775 |
| B_cells | Go_Positive_Regulation_Of_B_Cell_Proliferation | 0.7713 |
| B_cells | Initial_Triggering_Of_Complement | 0.7692 |
| B_cells | Go_Regulation_Of_Intrinsic_Apoptotic_Signaling_Pathway_By_P53_Class_Mediator | 0.7684 |
| B_cells | Go_Regulation_Of_Antigen_Processing_And_Presentation | 0.7679 |
| B_cells | Go_Regulation_Of_Intrinsic_Apoptotic_Signaling_Pathway_In_Response_To_Dna_Damage_By_P53_Class_Mediator | 0.765 |
| B_cells | Complement_Cascade | 0.7634 |
| B_cells | Go_Positive_Regulation_Of_Response_To_Cytokine_Stimulus | 0.7608 |
| B_cells | Go_Prostanoid_Metabolic_Process | 0.7606 |
| B_cells | Go_Regulation_Of_Dna_Damage_Response_Signal_Transduction_By_P53_Class_Mediator | 0.7604 |
| B_cells | Go_Negative_Regulation_Of_Intrinsic_Apoptotic_Signaling_Pathway_By_P53_Class_Mediator | 0.7603 |
| B_cells | Go_Cell_Recognition | 0.7553 |
| B_cells | Go_Regulation_Of_Macrophage_Cytokine_Production | 0.7542 |
| B_cells | Go_Mhc_Class_Ii_Protein_Complex | 0.7542 |
| B_cells | Go_Negative_Regulation_Of_Signal_Transduction_By_P53_Class_Mediator | 0.7503 |

**Supplementary Table 1.** MSigDB signatures most correlated (Pearson) with "B cell" signatures.

| Software | high throughput | comput-time | t-SNE interactive display | ease of use |
|---|---|---|---|---|
| Seurat | No | N/A | No display | + |
| ssGSEA | No | 4857 sec (*) | No display | + |
| AUCell | Yes | 1087 sec | Yes but not interactive | + |
| Single Cell Signature Explorer | Yes | 35 sec | Yes | +++ |

**Supplementary Table 2.** Summary of software comparison. Computation time to compute the scores for all KEGG database on Linux Xubuntu 18.10 workstation with two processors Xeon E5-2687w-v3 and 128Go RAM. (*) time extrapolated from 2 gene sets to whole KEGG

| S genes |
| --- |
| MCM5 |
| PCNA |
| TYMS |
| FEN1 |
| MCM2 |
| MCM4 |
| RRM1 |
| UNG |
| GINS2 |
| MCM6 |
| CDCA7 |
| DTL |
| PRIM1 |
| UHRF1 |
| MLF1IP |
| HELLS |
| RFC2 |
| RPA2 |
| NASP |
| RAD51AP1 |
| GMNN |
| WDR76 |
| SLBP |
| CCNE2 |
| UBR7 |
| POLD3 |
| MSH2 |
| ATAD2 |
| RAD51 |
| RRM2 |
| CDC45 |
| CDC6 |
| EXO1 |
| TIPIN |
| DSCC1 |
| BLM |
| CASP8AP2 |
| USP1 |
| CLSPN |
| POLA1 |
| CHAF1B |
| BRIP1 |
| E2F8 |

**Supplementary Table 3.** Genes used to define the S-phase

| G2M genes |
|---|
| HMGB2 |
| CDK1 |
| NUSAP1 |
| UBE2C |
| BIRC5 |
| TPX2 |
| TOP2A |
| NDC80 |
| CKS2 |
| NUF2 |
| CKS1B |
| MKI67 |
| TMPO |
| CENPF |
| TACC3 |
| FAM64A |
| SMC4 |
| CCNB2 |
| CKAP2L |
| CKAP2 |
| AURKB |
| BUB1 |
| KIF11 |
| ANP32E |
| TUBB4B |
| GTSE1 |
| KIF20B |
| HJURP |
| CDCA3 |
| HN1 |
| CDC20 |
| TTK |
| CDC25C |
| KIF2C |
| RANGAP1 |
| NCAPD2 |
| DLGAP5 |
| CDCA2 |
| CDCA8 |
| ECT2 |
| KIF23 |
| HMMR |
| AURKA |
| PSRC1 |
| ANLN |
| LBR |
| CKAP5 |
| CENPE |
| CTCF |
| NEK2 |
| G2E3 |
| GAS2L3 |
| CBX5 |
| CENPA |

**Supplementary Table 4.** Genes used to define the G2/M-phase

| adipocytes s | B cells | CD4 T cells | CD8 T cells | endothelial cells | $\gamma\delta$ T cells | myeloid cells | neuron | NK | pericytes | plasma cells |
|---|---|---|---|---|---|---|---|---|---|---|
| HBB | MS4A1 | TRBC1 | CXCR4 | FN1 | CXCR4 | LYZ | DKK3 | PRF1 | COL3A1 | IGHG3 |
| CFD | CD79A | TRBV19 | TRBC1 | EFEMP1 | AC092580.4 | S100A8 | PEG3 | GNLY | COL1A2 | CYAT1 |
| TNS1 | CD19 | CXCR4 | TRBV19 | COL4A2 | CCL5 | S100A9 | VSNL1 | NKG7 | SPARCL1 | IGLC1 |
| PLIN1 | CD79B | TRBC2 | PRF1 | CTGF | CCR5 | HLA-DRA | SLC1A2 | TRDC | CALD1 | IGKC |
| HBA1 | | TRBV6-5 | CD8A | SERPINE1 | CD247 | TYROBP | SPARCL1 | GZMB | RGS5 | IGHM |
| HBA2 | | TRBV5-4 | TRBC2 | THBS1 | CD3D | FCN1 | KIAA1211L | IL2RB | COL4A1 | IGK |
| GPX3 | | TRBV7-2 | TRBV6-5 | DKK3 | CD3G | CSTA | SYT1 | CCL5 | COL6A3 | IGHA2 |
| G0S2 | | TRBV3-1 | TRBV5-4 | COL8A1 | CD69 | CD14 | DCLK1 | CD247 | COL4A2 | IGLC7 |
| FABP4 | | CD52 | TRBV7-2 | COL4A1 | CREM | FGL2 | SYT4 | FGFBP2 | CYR61 | IGLC3 |
| LPL | | XIST | TRBV3-1 | TM4SF1 | CRIP1 | MNDA | CELF5 | GZMA | ATF3 | IGLC2 |
| RBP4 | | PTPRC | IL7R | CAV1 | CST7 | FCER1G | BEX1 | CX3CR1 | PRRX1 | IGLJ3 |
| CD36 | | TRAV20 | NKG7 | POSTN | EEF1G | MPEG1 | DNAJC6 | KLRF1 | FN1 | IGLJ2 |
| ADIPOQ | | TRDV2 | CD52 | GJA1 | FAM46C | SERPINA1 | DDAH1 | TYROBP | LAMC1 | IGLL5 |
| COL3A1 | | TRAJ17 | GZMH | FSTL1 | FGFBP2 | S100A12 | NEFM | CST7 | FERMT2 | FAM46C |
| LEP | | CD3D | GZMA | CALD1 | GNLY | CD52 | BEX5 | HOPX | FSTL1 | XIST |
| SEPP1 | | IL7R | PTPRC | LOXL2 | GPR171 | PLBD1 | SCG5 | KLRB1 | ADAMTS1 | JCHAIN |
| CIDEC | | LEF1 | CD3D | MGP | GZMA | CYBB | NEFL | CD300A | EGFL6 | TNFRSF17 |
| CRYAB | | CD2 | KLRK1 | CD93 | GZMB | NCF2 | MAP2 | TRBC1 | CAV1 | POU2AF1 |
| CHRDL1 | | IL2RB | KLRC4-KLRK1 | PODXL | GZMH | LST1 | NRCAM | TRBV19 | GEM | CXCR4 |
| CLEC3B | | TRAC | LCK | CYR61 | GZMK | ALOX5 | PNMAL1 | ADGRG1 | CTGF | MZB1 |
| SPARCL1 | | P2RY8 | CCL5 | SRPX | HOPX | TNFSF13B | PPP2R2C | TRDV3 | GPM6B | IGHD |
| CXCL14 | | ITK | TRAC | CDH5 | IL18RAP | RNASE6 | SMIM10L2A | CXCR4 | COL5A2 | TPD52 |
| ADIRF | | CD69 | TRAF3IP3 | SERPINE2 | IL2RB | FPR1 | SMIM10L2B | GZMH | DKK3 | GUSBP11 |
| DGAT2 | | CYTIP | TRGV9 | FRMD6 | IL32 | CPVL | PTPN5 | KLRK1 | TNS1 | IGHG2 |
| IGFBP5 | | TRBC1 | TRGC2 | PLS3 | IL7R | PTPRC | NEFH | KLRC4-KLRK1 | POSTN | IGHJ5 |
| PLIN4 | | TRBV19 | TRGC1 | LAMC1 | KLRB1 | MS4A6A | GRIA2 | RUNX3 | PLS3 | SLAMF7 |
| FSTL1 | | CXCR4 | TARP | FHL2 | KLRC1 | CD93 | TPD52 | PTPRC | IGFBP5 | RGS1 |
| COL4A2 | | TRBC2 | P2RY8 | TGM2 | KLRC2 | CXCR4 | PTPRD | PLAC8 | RND3 | IGLV@ |
| CXCL12 | | TRBV6-5 | FGFBP2 | WBP5 | KLRC4-KLRK1 | ITGAM | MYT1L | IL18RAP | MATN2 | CCR2 |
| PCOLCE2 | | TRBV5-4 | PLAC8 | DDAH1 | KLRD1 | HCK | TCEAL2 | CCL4 | EPS8 | IGHA1 |
| C10orf10 | | TRBV7-2 | GNLY | PTPRK | KLRK1 | CD36 | PTPRK | TRBC2 | KITLG | IGHG1 |
| COL1A2 | | TRBV3-1 | LAT | MMP2 | LCK | | FAT3 | TRBV6-5 | FBN1 | IGHJ4 |
| PPP1R1A | | CD52 | RUNX3 | TFPI | MIAT | | GPR158 | TRBV5-4 | EDNRA | ZCWPW2 |
| CALD1 | | XIST | CD2 | PTX3 | MT2A | | RGS4 | TRBV7-2 | COL12A1 | DERL3 |
| FERMT2 | | PTPRC | GZMK | | NKG7 | | ELAVL2 | TRBV3-1 | TM4SF1 | CD38 |
| COL6A3 | | TRAV20 | CST7 | | P2RY8 | | NELL2 | SH2D1B | GPNMB | CTHRC1 |
| AOC3 | | TRDV2 | CD69 | | PRF1 | | DOCK4 | MYBL1 | AOC3 | AMPD1 |
| CAV1 | | TRAJ17 | BCL11B | | PTPRC | | GRM3 | KLRC1 | C8orf4 | SPAG4 |
| LOC102724852 | | CD3D | | | RGS1 | | SEPP1 | KLRC2 | TJP1 | CSF2RB |
| H19 | | IL7R | | | RORA | | TENM2 | KLRD1 | LUM | TBC1D9 |
| LAMC1 | | LEF1 | | | RPL17 | | DIRAS2 | CTSW | ECM2 | EAF2 |
| COL4A1 | | CD2 | | | RPL36A | | ELAVL4 | ITGAM | FHL2 | CD27 |
| SAA1 | | IL2RB | | | RPS26 | | GPM6B | CD52 | ACTG2 | ANKRD36BP2 |
| SAA2 | | TRAC | | | RUNX3 | | PCDH7 | LCK | WBP5 | PIP5K1B |
| EFEMP1 | | P2RY8 | | | SLC7A5 | | STMN2 | XCL1 | SLC7A2 | HLA-DOB |
| VWF | | ITK | | | TARP | | RORB | FCER1G | NR2F2 | LAX1 |
| MAOB | | CD69 | | | TRDC | | SH3GL2 | S1PR5 | FGL2 | IGLL3P |
| GYG2 | | | | | TRDV3 | | CNTN3 | XCL2 | | |
| | | | | | TRGC1 | | CNDP1 | | | |
| | | | | | TRGC2 | | | | | |
| | | | | | TRGV9 | | | | | |
| | | | | | XIST | | | | | |

**Supplementary Table 5.** Genes used in supplementary figure 2

| normalization | computation time |
|---|---|
| raw | 0m 23s |
| FTMN | 8m 44s |
| Seurat | 0m 22s |

**Supplementary Table 6.** Elapsed computing time to calculate the signature enrichment scores of all gene sets from KEGG database by Single-Cell Signature Scorer for the 8k PBMC dataset using either the raw UMI data, FTMN noise-reduced UMI, or the Seurat- normalized UMI data.
